# Insertions in SARS-CoV-2 genome caused by template switch and duplications give rise to new variants that merit monitoring

**DOI:** 10.1101/2021.04.23.441209

**Authors:** Sofya K. Garushyants, Igor B. Rogozin, Eugene V. Koonin

## Abstract

The appearance of multiple new SARS-CoV-2 variants during the winter of 2020-2021 is a matter of grave concern. Some of these new variants, such as B.1.617.2, B.1.1.7, and B.1.351, manifest higher infectivity and virulence than the earlier SARS-CoV-2 variants, with potential dramatic effects on the course of the COVID-19 pandemic. So far, analysis of new SARS-CoV-2 variants focused primarily on point nucleotide substitutions and short deletions that are readily identifiable by comparison to consensus genome sequences. In contrast, insertions have largely escaped the attention of researchers although the furin site insert in the spike protein is thought to be a determinant of SARS-CoV-2 virulence and other inserts might have contributed to coronavirus pathogenicity as well. Here, we investigate insertions in SARS-CoV-2 genomes and identify 347 unique inserts of different lengths. We present evidence that these inserts reflect actual virus variance rather than sequencing errors. Two principal mechanisms appear to account for the inserts in the SARS-CoV-2 genomes, polymerase slippage and template switch that might be associated with the synthesis of subgenomic RNAs. We show that inserts in the Spike glycoprotein can affect its antigenic properties and thus merit monitoring. At least, three inserts in the N-terminal domain of the Spike (ins245IME, ins246DSWG, and ins248SSLT) that were first detected in 2021 are predicted to lead to escape from neutralizing antibodies, whereas other inserts might result in escape from T-cell immunity.

## Main text

The first SARS-CoV-2 genome was sequenced in January 2020. Since then, more than a milion virus genomes have been collected and sequenced. Comparative analysis of SARS-CoV-2 variants has provided for the identification of the routes of virus transmission ^1–4^, the selective pressure on different genes ^5^, and the discovery of new variants associated with higher infectivity ^6–8^. In many cases, genome analysis only included search for point mutations, but some deletions also have been identified, such as del69-70, one of the characteristic mutations of B.1.1.7 and Cluster 5 ^2,3^ or del157-158 in B.1.617.2 (delta) ^9^. Moreover, recently, recurrent deletions have been shown to drive antibody escape ^10^. However, insertions are mostly ignored, both during variant calling step and in the downstream analysis.

Although insufficiently studied, insertions appear to be crucial for beta-coronavirus evolution. Three insertions in the spike (S) glycoprotein and in the nucleoprotein (N), that occurred early in sarbecovirus evolution, have been shown to differentiate highly pathogenic beta-coronaviruses (SARS-CoV-1, SARS-CoV-2 and MERS) from mildly pathogenic and non-pathogenic strains, and suggested to be key determinants of human coronaviruses pathogenicity ^11^. The best characterized insert in SARS-CoV-2 is the PRRA tetrapeptide that so far is unique to SARS-CoV-2 and introduces a polybasic furin cleavage site into the S protein, enhancing its binding to the receptor ^12–14^.

Inserts in the SARS-CoV-2 genome are categorized in the CoV-GLUE database ^15^, and the preliminary results on systematic characterization of the structural variance and inserts in particular have been reported ^16^. Forty structural variants including three inserts, three nucleotides long each, were discovered and shown to occur in specific regions of the SARS-CoV-2 genome. These variants have been further demonstrated to be enriched near the 5’ and 3’ breakpoints of the non-canonical (nc) subgenomic (sg) RNAs of coronaviruses. Additionally, indels have been shown to occur in arms of the folded SARS-CoV-2 genomic RNA ^16^. However, longer inserts that might have been introduced into the virus genome during SARS-CoV-2 evolution, to our knowledge, have not been systematically analyzed.

The mechanisms of sequence insertion in the genomes of RNA viruses, and coronaviruses in particular, are poorly understood. One potential route is recombination. Homologous recombination is common among coronaviruses, and in particular, in the sarbecovirus lineage, and is likely to be a major evolutionary route producing coronavirus strains with changed properties ^17,18^. Specifically, the entire receptor-binding motif (RBM) domain of the S protein can be replaced by homologous recombination as it probably happened in RaTG13 and some other sarbecoviruses ^17,19,20^. In contrast, non-homologous recombination in RNA viruses appears to be rare, and its molecular mechanisms remains poorly understood ^21^.

In infected cells, beta-coronaviruses produce 5 to 8 major sgRNAs ^22,23^. Eight canonical sgRNAs are required for the expression of all encoded proteins of SARS-CoV-2. These sgRNAs are produced by joining the transcript of the 5’ end of the genome (TRS site) with the beginning of the transcripts of the respective open reading frames (ORFs) ^24^. In addition, SARS-CoV-2 has been reported to produce multiple nc sgRNAs, some of which include the TRS at 5’ end, whereas others are TRS-independent ^25,26^; apparently, the ncRNAs are spurious products of errors of transcription initiation.

Here we report the comprehensive census of the inserts that were incorporated into virus genomes during the evolution of SARS-CoV-2 over the course of the pandemic and show that occurred during virus evolution rather than resulting from experimental errors. These inserts are non-randomly distributed along the genome, most being located in the 3’terminal half of the genome and co-localizing with 3’ breakpoints of nc-sgRNAs. We show that the long insertions occur either as a result of the formation of nc-sgRNAs or by duplication of adjacent sequences. We analyze in detail the inserts in the S glycoprotein and show that at least two of these are located in a close proximity to the antibody-binding site in the N-terminal domain (NTD), whereas others are also located in NTD loops and might lead to antibody escape, and/or T cell evasion.

## Results

### Identification of inserts in SARS-CoV-2 genomes

To compile a reliable catalogue of inserts in SARS-CoV-2 genome, we analyzed all the 1,785,103 sequences present in the GISAID multiple genome alignment (compiled on June 17, 2021). From this alignment, we extracted all sequences that contained insertions in comparison with the reference genome (1354 unique events in 2159 unique genomes). After the initial filtering (Materials and Methods), insertions were identified in 752 unique genomes, with 544 unique events detected in total. We evaluated all regions around insertions in alignments and removed all inserts that appeared due to misalignment. To the remaining inserts, we added four long inserts obtained from the GISAID metadata description (see Materials and Methods), resulting in a set of 354 unique inserts ranging in length from 2 to 69 nucleotides in 746 genomes (including identical sequences) (Figure 1a; Supplementary Table 1; Supplementary Table 2).

**Figure 1.**
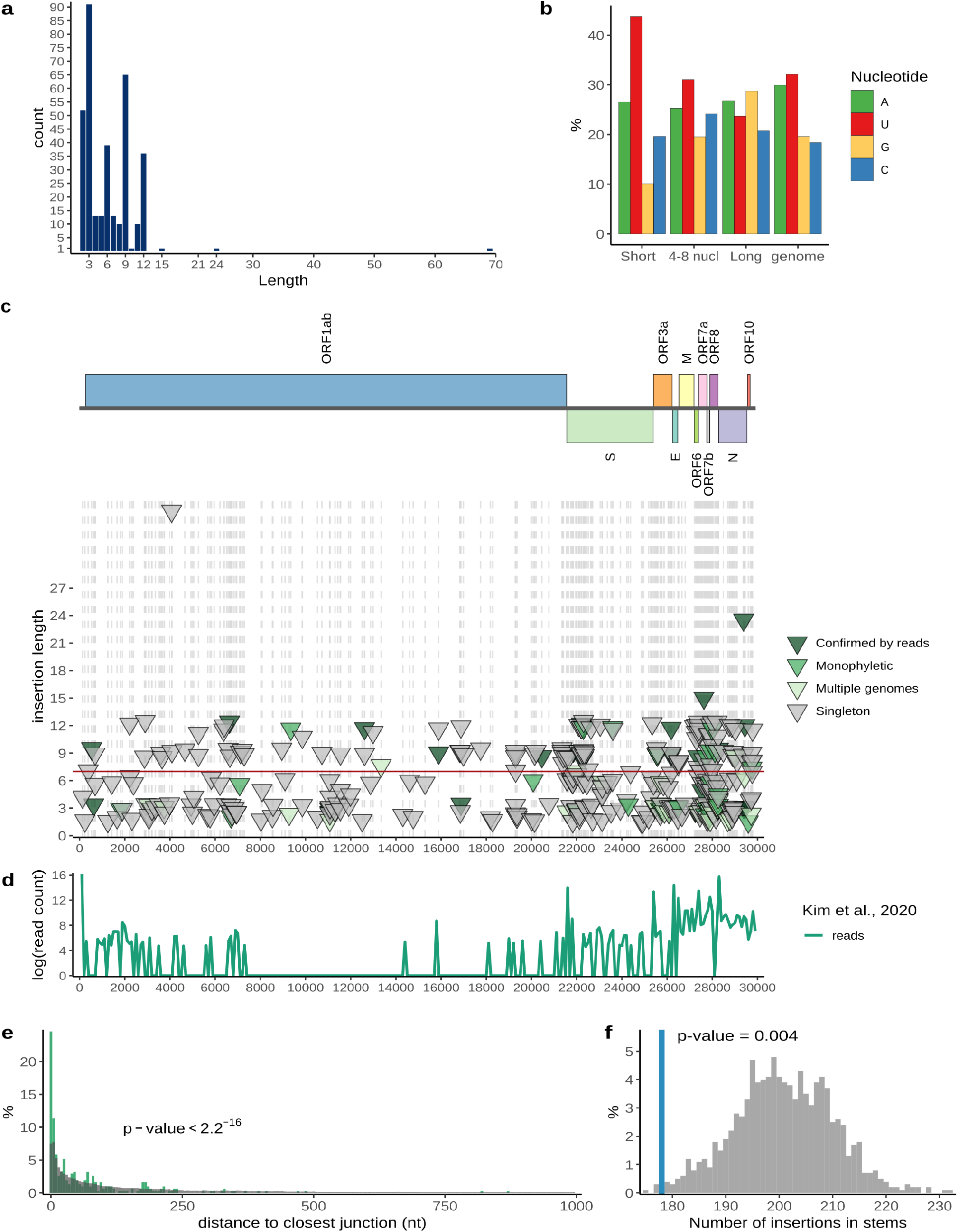
Insertions in SARS-CoV-2 genomes. (a) Distribution of insert lengths. (b) Nucleotide composition of inserts of different lengths and full SARS-CoV-2 genome. (c) Distribution of inserts along the genome. Each triangle represents one insertion event. The level of confidence in each variant is represented by color: dark green, confirmed by sequencing read analysis; green, monophyletic in the tree, no read data available; light green, observed multiple times, but not monophyletic; grey, singletons (Supplementary Table 2). The positions of inserts are marked with grey dashed lines. (d) Experimental data on SARS-CoV-2 transcriptome ^26^ showing template switch hotspots during the formation of sgRNAs, showing the distribution of junction reads connecting recombination hotspots along the genome. (e) Distance from inserts to closest template switching hotspot site (green) compared with random expectation (grey). Wilcoxon rank sum test p-value is provided. (f) The number of inserts that occur in structured regions of SARS-CoV-2 genomic RNA (blue) compared with random expectation (grey). Permutation test p-value is provided. The data on SARS-CoV-2 structure was obtained from ^28^.

To further minimize the number of inserts that appeared due to sequencing errors, we screened the Sequence Read Archive (SRA) database for the corresponding raw read data. We were able to obtain raw reads for 43 inserts, of which 40 were multiples of three by length, and one more was four nucleotides but occurred in an intergenic region. We validated 35 insertions with raw read data; only one of these was not a multiple of three in length and occurred within a gene (Supplementary Table 1; Supplementary Table 2). Among the inserts that were not validated by raw reads 6 were singletons, whereas two others were short duplets and occurred in a polyU tract. We removed these unconfirmed events from our dataset, resulting in 346 unique inserts. Assuming that the fraction of true positives is the same among all inserts as it is among those with available reads, 282 of these inserts are expected to reflect actual evolutionary events. Among the inserts in our dataset, 234 (67%) were multiples of three, and of the remaining ones, 16 (5% of the total) were located in intergenic regions, 39 (11%) in orf1ab, and 57 (17%) in other genes. It appears likely that most if not all frameshifting inserts are sequencing artifacts, but some of such inserts in other genes could be real events reflecting the dispensability of these genes for virus reproduction. For example, we identified 4 frameshifting inserts in ORF6, for which deletion variants have been described ^27^

The short inserts (< 9 nt) had a distinct nucleotide composition with a substantial excess of uracil, at about 45%, whereas the composition of the long inserts (9 nt or longer) was similar to that of the SARS-CoV-2 genome average, with about 25% U (Figure 1b). This trend was even more pronounced for inserts verified by raw reads (Supplementary Figure 1). Thus, we split the collection of inserts into two categories, the short and long inserts, which we analyzed separately.

We then checked whether inserts that were present in multiple genome sequences were located close to each other in the global phylogenetic tree (see Materials and Methods). We did not require strict monophyly because inserts are not always included in the genome sequences (see Supplementary note 1). Of the 52 short inserts identified in multiple genomes, 18 met this relaxed monophyly criterion (Supplementary Table 2). In 5 cases, identical insertions were observed in genomes submitted from the same laboratory, and mostly, on the same date, which implies that the genomes were sequenced and analyzed together, making it difficult to rule out a sequencing error. Interestingly, 7 of the 8 cases that were validated by raw reads were not monophyletic. By contrast, among the 15 long inserts that were found in multiple genomes, 14 met the monophyly criteria including all 5 validated by raw reads (Supplementary Table 2).

In summary, the inserts detected in SARS-CoV-2 genomes fell into the following categories: 230 (66.5%) short inserts, among which 15 (4.3% of the total) were validated by raw reads; and 116 long (at least, 9 nt) (33.5%) inserts. We additionally classified the long inserts into four groups, in the order of increasing confidence: 87 (25%) singletons, 1 non-monophyletic insert observed in two genomes, 9 (2.6%) monophyletic inserts observed in multiple genomes, all with no available raw reads and 20 (5.8%) inserts (15 singletons and 5 monophyletic ones), for which the insertions were validated by the raw reads. The 15 (4.3%) short inserts confirmed by read data and 29 (8.4%) long inserts that were detected in multiple genomes (monophyletic and not) and/or confirmed by raw reads comprised the set of the most reliable insertion events that were observable across the evolution of SARS-CoV-2 during the pandemic (Supplementary Table 4). Among these highly reliable inserts, there was only one frameshifting insert within orf6 gene.

### Insertions are non-uniformly distributed along the SARS-CoV-2 genome

We found that the insertions were not randomly distributed along the genome, with most occurring in the 3’-terminal third of the genome (Figure 1c). Two, not necessarily mutually exclusive main hypotheses have been proposed on the origin of the short inserts (structural variants) in coronavirus, namely, that they are associated with loops in the virus RNA structure or occur near the hotspots of template switch, at the breakpoints of TRS-independent transcripts ^16^. To differentiate between these two mechanisms, we compared the distribution of 347 inserts along the SARS-CoV-2 genome with the distributions of structured regions ^28^ and of template switch hotspots ^26^. We detected a strong association of the insertions with the template switch hotspots (Pearson correlation r = 0.42, p-value = 1.8×10^−14^) (Figure 1d). Almost 25% of the inserts occurred within 5 nt of a template switch hotspot, compared to less than 10% expected by chance, and the distribution of lengths observed in the real data is significantly different from random expectation (Wilcoxon test p-value < 2.2×10^−16^; Figure 1e). Also, we observed that inserts were significantly less frequent in predicted RNA stems than expected by chance (permutation test p-value = 0.004; Figure 1f). Both these observations held, with statistical significance, when we analyzed only the 45 highly confident insert (Supplementary Figure 4). Thus, inserts in SARS-CoV-2 genomes are associated with template switch hotspots, and also tend to occur in RNA loops.

### Short insertions in SARS-CoV-2 are generated by template sliding

The notable difference in nucleotide composition and different phyletic patterns of short and long inserts imply that the two types of insertions occur via different mechanisms. As pointed out above, the short insertions are rarely monophyletic, indicating that short U-rich sequences were inserted in the same position in the SARS-CoV-2 genome on multiple, independent occasions during virus evolution in the course of the pandemic. Also, 43 of the 230 short inserts occurred in runs of U or A, whereas 63 more probably result from local duplications (Supplementary table 2). These observations suggest that short insertions occur via template sliding (polymerase stuttering) on runs of As or Us in the template (negative strand or positive strand, respectively) RNA ^29–31^ (Supplementary figure 5a; Supplementary table 2). This could be either a biological phenomenon occurring during SARS-CoV-2 evolution, in case the errors are produced by stuttering of the coronavirus RdRP, or an artifact if the errors come from the reverse transcriptase or DNA polymerase that is used for RNA sequencing, or are a mix of biological and experimental polymerase errors. However, for all 17 short inserts that were confirmed by raw read analysis, we also detected the U enrichment (Supplementary Figure 1). Those inserts were observed at high allele frequencies in the data (Supplementary Table 1), and thus, are unlikely to result from experimental errors. Additionally, short inserts appear to be represented with the same frequency in SARS-CoV-2 genomes sequenced with different technologies, including Illumina MiSeq, NovoSeq and NextSeq, and even Oxford Nanopore or IonTorrent (Supplementary Table 1). Furthermore, elevated rate of thymine insertion has not been reported as a common error of either Illumina or Oxford Nanopore technology ^32–35^. In contrast, production of longer transcripts and slow processing on polyU tracts has been demonstrated for nsp12 (RdRP) of SARS-CoV-1 ^36^. Additionally, the RdRp complex of SARS-CoV lacking the proof-reading domain has been shown to misincorporate more nucleotides compared with other viral polymerases ^37^. Thus, it appears likely that most of the short inserts in SARS-CoV-2 genomes are generated by stuttering of the virus RdRP.

### Long insertions in SARS-CoV-2 are caused by template switching and local duplications

For in-depth analysis of the long inserts, we selected only the 29 high-confidence ones (see above), which were found in 74 genomes and ranged in size from 9 to 24 nucleotides (Figure 2, Supplementary Table 4).

**Figure 2.**
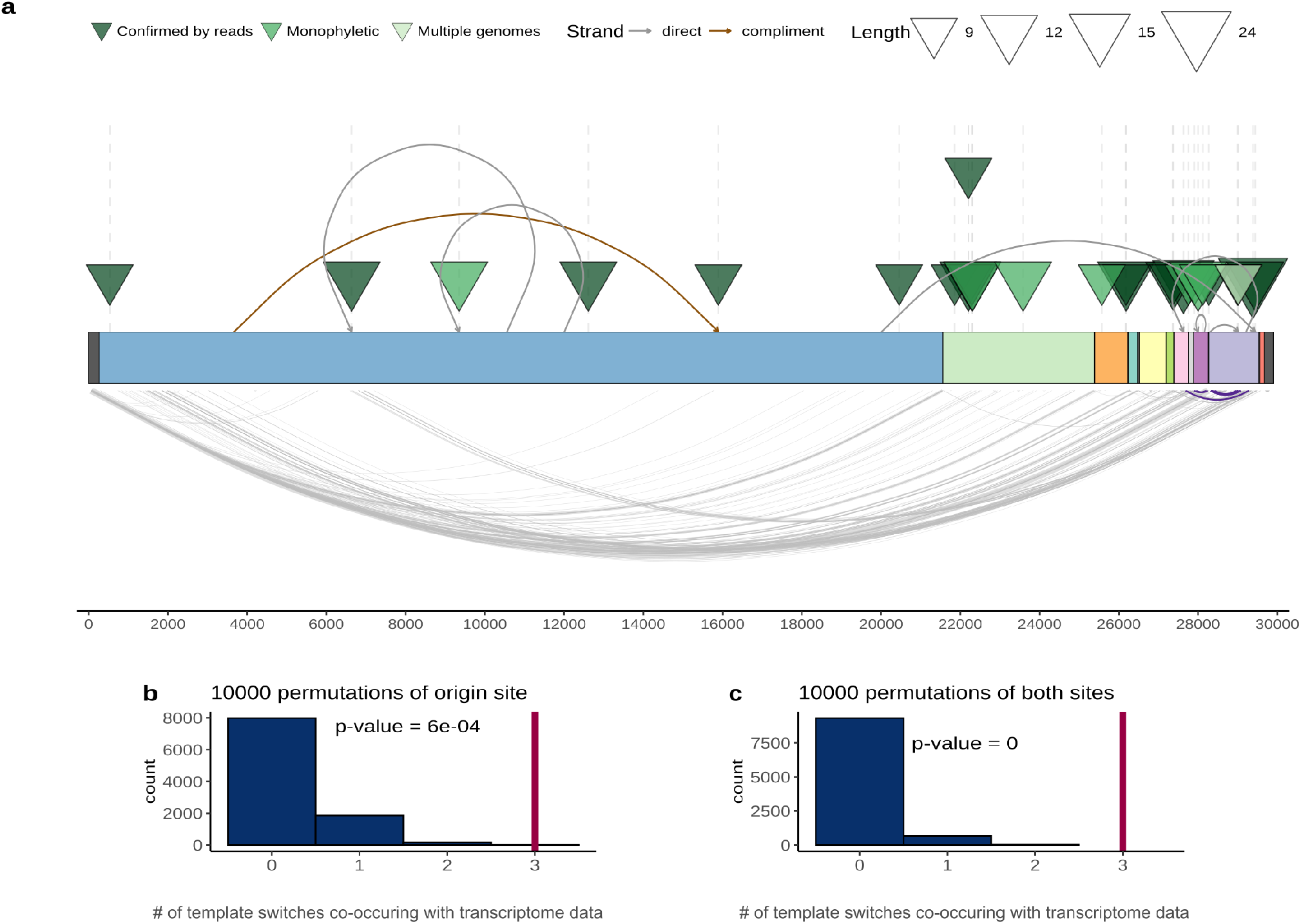
Long insertions possibly occur through template switching and formation of nc sgRNAs. (a) Each triangle shows an independent insertion event, colored as in Fig. 1. Curves on the upper side of the plot connect the insertion origin site and insertion position, brown color indicates that the origin sequence is on the same strand, and grey color shows that the origin sequence is on complementary strand, Curves at the bottom of the plot represent the experimental data on sgRNAs from Kim et al. ^26^. Curves highlighted in violet correspond to the three cases when insert and corresponding origin site co-occur with sgRNA junctions. The SARS-CoV-2 genes are colored as in Fig.1. Permutation tests show the number of template switches co-occurring with RdRp jumps (x-axis) expected at random (blue), (b) when only the positions of the origins were randomly sampled 10000 times from the genome. (c) when both ends were randomly sampled. Red vertical line represents the number observed in data.

Those insertions originated from different laboratories that employed different protocols. Furthermore, these events started to accumulate in late April 2020, and the median collection date of the genomes containing the long inserts is February, 8 2021. Eight of the 29 reliable long insertions are located in the S gene, which is significantly more than expected by chance (Binomial test p-value = 0.027). The excess of inserts in the S gene suggests that their spread in the virus population could be caused by positive selection, perhaps, driven by enhancement of the interactions of SARS-CoV-2 with the host cells conferred by the inserts. This possibility is in agreement with the recent detection of an excess of amino acid replacements apparently due to positive selection in the S gene ^38^, although a contribution of relaxed purifying selection cannot be ruled out either.

The lengths of these high-confidence inserts allowed us to search for matching sequences both in SARS-CoV-2 genomes and in other viruses. For 9 cases, we were able to identify the probable origin of the insertion. All detected matches were within the SARS-CoV-2 genome and contained no substitutions. For two inserts, we detected a local duplication that most likely gave rise to the insertion (Supplementary Figure 5b). In both cases, these inserts were found in multiple genomes and are probably monophyletic, and one of these inserts was validated with raw read data. In 7 more cases, we detected significant matches in the SARS-CoV-2 genome, 6 in the coding strand and one in the complementary strand (Figure 2a; Supplementary Table 4). Among these 7 insertions, two were monophyletic, and four more were singletons supported by raw data. The apparent origin of inserts from distant parts of the SARS-CoV-2 genomes implies template switch (Supplementary Figure 5c). In three more cases, we were able to propose possible sources of inserts in SARS-CoV-2 genome, but the matches did not reach statistical significance (Supplementary Table 4). Although we could not find a probable source of many inserts, which is a limitation of our analysis, the principal reason of this failure is that most inserts are only 9 nt long, so that there is not enough statistical power for detecting likely origin sequences with substitutions.

We hypothesized that the long inserts, at least those, for which potential origin sequences were detected, originate from template switching that occurs during the synthesis of the nc sgRNAs, when the RdRP jumps from one genome location to another. To assess the possibility that RdRP jumping contributes to insertions, we compared the insert locations and the sites of the likely origin of the inserts with the experimental data on the SARS-CoV-2 transcriptome ^26^. The jumps produce junction reads that connect two distant locations in the genome. Regions that are connected by multiple reads are hotspots of template switching (Figure 2). If the insertion sequence and the origin sequence are located close to these junction sites, it is likely that they were introduced by the RdRP during sgRNA synthesis. As pointed out above, inserts show a non-random proximity to template switch hotspots, so for the inserts with a traceable origin, we additionally checked whether their sites of origin occurred close to the sites of RdRp jumping. Although the information on the SARS-CoV-2 transcriptome is limited, among the 7 inserts with predicted origins, we found that three insert sites were located within one end of the junction, whereas their corresponding sites of origin were within 100 nucleotides of the other side of the same junction (Figure 2a). To assess the significance of this finding, we performed two permutation tests (see Material and Methods), in one of which the real insertion positions were matched with start sites chosen randomly, whereas in the second one, both types of sites were selected at random. Both tests showed significant co-localization of the inserts with template switch junctions (Figure 2 b,c).

Thus, high-confidence long inserts in the SARS-CoV-2 genome probably originated either by local duplication or by template switching which, at least in some cases, seemed to be associated with nc sgRNA synthesis. Notably, the PRRA insert, the furin cleavage site that is one of characteristic features of SARS-CoV-2, resembles the long inserts analyzed here. Although this insert has a high GC-content compared to the genomic average of SARS-CoV-2, it falls within the GC-content range of the long inserts (Supplementary Figure 1b). Furthermore, this insert is located within 20 nucleotides of a template switch hotspot at position 22,582 ^26^. Although we were unable to identify a statistically significant match that would allow us to map the origin of the PRRA insert to a particular location within the SARS-CoV-2 genome, this insert also might have originated by template switch, with subsequent substitutions erasing the similarity to the origin sequence.

### Insertions in the S protein produce putative antibody escape variants

As indicated above, insertions are non-uniformly distributed along the SARS-CoV-2 genome (Figure 1c). In particular, among the 29 long inserts identified with high confidence, 8 were located in the S protein, suggesting that these inserts could persist due to their adaptive value to the virus. Four of the 8 inserts in S were observed in multiple genomes that formed compact clades in the phylogenetic tree (Supplementary note 1), and two (ins214AAG and ins214TDR) were strongly supported by raw reads. Four more inserts (ins98KAE, ins214KLGP, ins245IME, and ins246DSWG) were found in single genomes, but again, were strongly supported by raw reads, where they reached allele frequency close to unity, so these are highly unlikely to be artifacts (Supplementary Table 1). Inserts in Spike were observed in various PANGO lineages, some of which are circulating to date, what is more ins246DSWG happen within B.1.1.7 (Alpha) lineage (Supplementary table 4; Supplementary note 2).

All 8 long inserts in S protein were located on the surface, within computationally predicted epitope candidates, and could potentially modify the interactions of S with the receptor and/or antibodies ^36^ (Figure 3a). Seven of these 8 eight inserts mapped to the N-terminal domain (NTD) of S, and three of these occurred in the same genome position, 22,004 (Figure 3). Compared to the receptor binding domain, the NTD initially attracted much less attention. Subsequently, however, multiple substitutions associated with variants of concern and observed in immunocompromised individuals with extended COVID-19 disease were identified in the NTD ^2,40,41^. To evaluate potential functional effects of the inserts in the NTD, we mapped them onto the protein structure. Three inserts, ins245IME, ins246DSWG and ins248SSLT, are located in the loop that is responsible for the interaction with the 4A8 antibody and potentially other antibodies ^42^ (Figure 3). Thus, at least these three insertions might be associated with the escape of SARS-CoV-2 variants from immune antibodies. The presence of multiple insertions in the same site 22,004 is suggestive of a role in infection, which is compatible with the observation of multiple deletion variants in the same region, in particular 21971-22005 ^10^. These insertions and ins98KAE are located in the neighboring loops, and given that the central region of the NTD is essential for the virus interaction with CD4+ cells ^43^, could be associated with the escape from the T-cell immunity. Also, there is additional evidence that this region could represent another epitope for antibody binding ^44^. Because these insertions were detected only in recent samples, it appears that the respective variants merit further monitoring although they have not reached a high frequency in the virus population.

**Figure 3.**
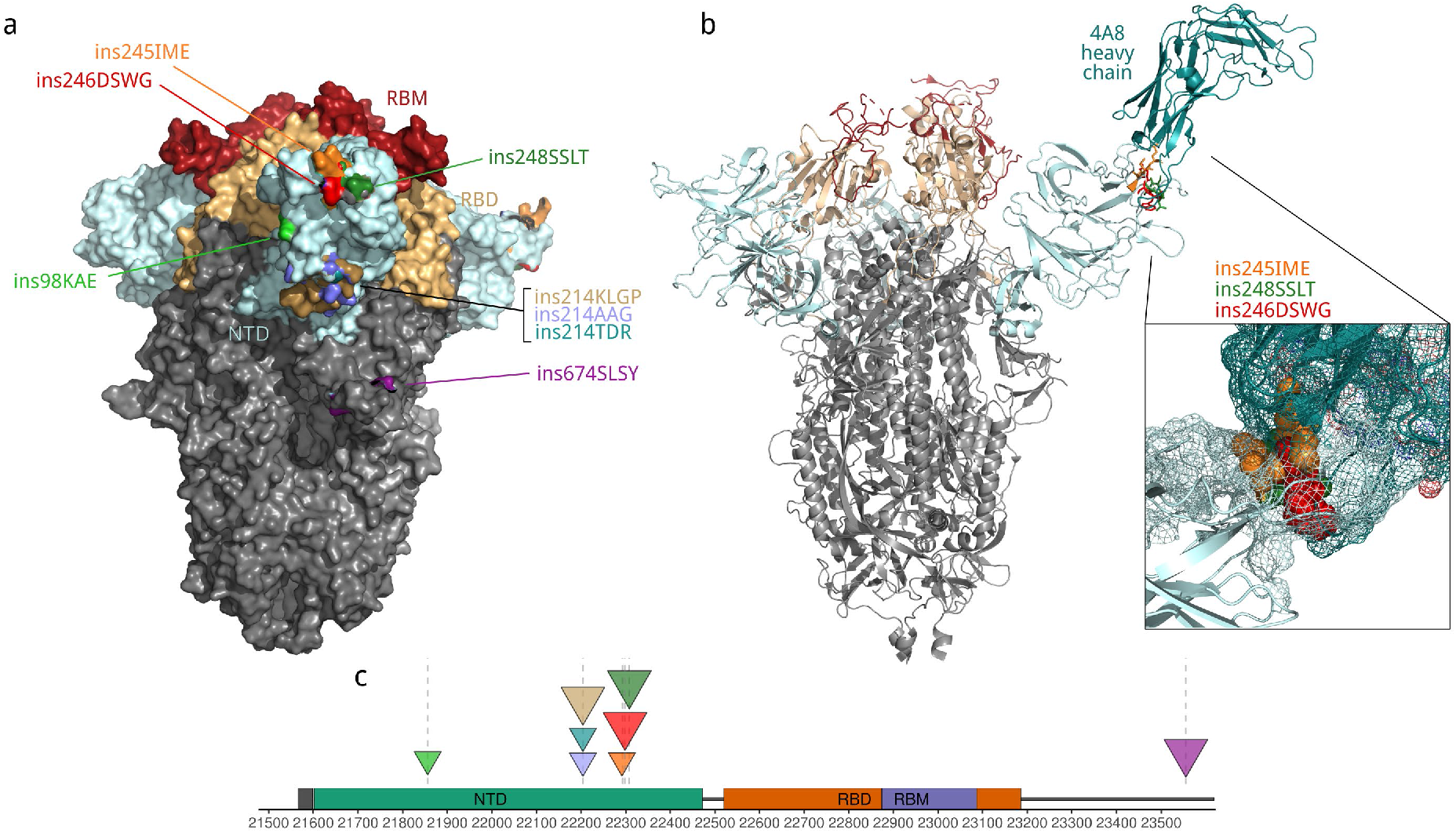
Location of insertion sites in SARS-CoV-2 S protein. (a) Surface representation showing that all observed insertions can potentially change surface properties (PDB ID: 7cn8). (b) Ins 245, 246 and 248 are located on the surface interacting with 4A8 antibody (PDB ID: 7cl2). Enlarged is the interacting surface. Cyan, N-terminal domain (NTD), wheat, receptor-binding domain (RBD), dark red, receptor binding motif (RBM), aquamarine, heavy chain of the 4A8 antibody (PDB ID: 7cl2). Each insertion is shown in a distinct color. The models for each insertion were generated with the SWISS-model web server. (c) Location of insertions in the genome of SARS-CoV-2. Full description of insertions is provided in the Supplementary Tables 4. Triangle size is proportional to the insert length.

## Discussion

Although structural variation is an important driver of betacoronaviruses evolution, in the genome analysis during the COVID-19 pandemics, part of the structural variations, namely, long insertions, to our knowledge, have not been systematically analyzed. This is a potentially consequential omission, given that insertions in the S and N proteins might contribute to the betacoronavirus pathogenicity. In particular, the furin cleavage site inserted into the S protein seems to be crucial for SARS-CoV-2 pathogenicity ^45,46^, and insertions that seem to differentiate high-pathogenicity coronaviruses from low pathogenicity ones have been detected in both S and N ^11^. Furthermore, betacoronaviruses are known to produce transcripts of greater than genome length ^24^, suggesting that insertions occur frequently during the reproduction of these viruses. Here we attempted a comprehensive identification and analysis of insertions in the SARS-CoV-2 protein-coding sequences that originated during the current pandemic.

We found that short and longer insertions substantially differed with respect to their nucleotide compositions and mapping to the phylogenetic tree, suggesting that different mechanisms could be at play. The short inserts were strongly enriched in U and in most cases occurred independently in the tree. It appears likely that these inserts resulted from RdRP slippage on short runs of A or U. Indeed, the observed excess of U in these inserts resembles the error profile of SARS-CoV-1 RdRP ^36^. In contrast, the composition of the long inserts was close to that of the virus genome, and many of these insertions were found to be monophyletic, that is, these appear to be rare events that did not occur at nucleotide runs. Sequence analysis of the SARS-CoV-2 genomes indicates that these insertions occur either through polymerase slippage resulting in tandem duplication or more commonly, seem to have been triggered by illegitimate template switching associated with the formation of nc sgRNAs. In support to our hypothesis template switching in different RNA viruses has been demonstrated previously in a variety of experimental settings ^47,48^. Furthermore, template switching has been observed in coronaviruses^49^.

For approximately one third of the long insertions, we were unable to pinpoint the source of the inserted sequence. This could be explained simply by the mutational deterioration of the similarity between the source and insert sequences, especially, for 9 nt inserts. However, a third mechanism of insertion cannot be ruled out. The PRRA insert that comprises the furin cleavage site in the S protein resembles the long inserts and likely originated by similar mechanisms although its origin by template switching and recombination with another sarbecovirus remains a possibility ^50^.

Long inserts are markedly overrepresented in the S glycoprotein, particularly, in the NTD. Examination of the locations of these inserts on S protein structure strongly suggests that at least some of the inserts in the NTD result in the escape of the respective variants from neutralizing antibodies and, possibly, also from the T-cell response. The excess of insertions in the S protein is compatible with this protein being the principal area of virus adaptation. However, the location of most of the inserts in the NTD, as opposed to the RBD, appears unexpected. Considering that all the detected inserts appeared at a relatively late stage of the pandemic, it seems likely that the structure of the RBD was already largely optimized for receptor binding at the onset of the pandemic such that most insertions would have a deleterious effect. In contrast, insertions into the NTD might allow virus to escape immunity without compromising the interaction with the host cells. Thus, the insertion variants appear to merit monitoring, especially, at a time when vaccination might select for escape variants.

## Materials and methods

### GISAID data

The full multiple alignment of 1785103 complete SARS-CoV-2 genomes (version 0617) was downloaded from GISAID (https://www.gisaid.org/). From this alignment, we extracted all positions of insertions. An insertion was defined as addition of any number of columns compared to the SARS-CoV-2 reference genome (hCoV-19/Wuhan/Hu-1/2019 (NC_045512.2)). All insertions detected in the first and last 100 positions of the reference sequence were discarded as potentially erroneous. The alignment around the potential insertions was manually inspected, and all inserts that were the result of misalignment were discarded. All the sequences that had more than two insertions were discarded, in order to avoid genomes with multiple sequencing errors. Also all inserts with ambiguous symbols were discarded. Information on the laboratory of origin, sequencing platform and consensus assembly methods (where available) was extracted from GISAID metadata. Additionally, because GISAID removes from alignment all inserts > 12 nucleotides that appeared only once, we downloaded the metadata for all genomes with inserts, that were submitted before 2021-06-24, and selected those events that were at least 12 nucleotides long. We retained only those that had corresponding data in SRA, and were confirmed by raw read data analysis (see below).

To identify all genomes that contained the particular insert, we downloaded all GISAID sequences available by 2021-06-23 and used cdhit-est-2d to find identical sequences with parameters: -s 0.99 -c 1.0 -n 11.

### Insertion validation from raw read data

Raw reads were downloaded from SRA database (https://www.ncbi.nlm.nih.gov/sra) with SRA Toolkit (Supplementary Table 1). The reads were mapped to the SARS-CoV-2 reference genome (NC_045512.2) with bowtie2 version 2.2.1 ^51^, either in pair mode of single read mode, depending on the type of data deposited to the SRA. The variants in each genome were called with LoFreq version 2.1.5 ^52^ as described in Galaxy (https://github.com/galaxyproject/SARS-CoV-2/blob/master/genomics/4-Variation/variation_analysis.ipynb). All insertions identified with LoFreq were visualized with the IGV software and manually inspected. An insertion was considered a real biological event if it had an allele frequency in reads more than 50%, was located in the middle of the amplification fragment, and was covered by at least 10 reads.

### Search for origins of long insertions

Search for putative duplications/template switch events with and without mismatches was performed against SARS-CoV-2 and closely related SARS-CoV genomes from human and 43 bats (https://hgdownload.soe.ucsc.edu/goldenPath/wuhCor1/multiz44way/). A pangolin SARS-CoV (MT040335.1) was added to this dataset. Each insertion sequence was compared to all subsequences from a target sequence. All sequences with either the perfect match or with mismatches was retrieved (putative insertion source, PIS). If a PIS was located immediately upstream or downstream of an insertion sequence, it was annotated as duplication. If the PIS was located in any other positions, the template switch model was accepted as the best explanation of the observed insertion sequence. This procedure was implemented using a sliding window, of a length equal to the length of the analyzed insert. If no perfect match was detected, the window with the minimal number of mismatches was retrieved and considered a putative insertion source.

To assess the significance of putative duplications and template switch events, we designed a sampling procedure to test a hypothesis that an insertion is not the result of spurious matches between an insertion sequence and corresponding PIS. Each insertion sequence was shuffled and scanned against datasets using the sliding window described above. This procedure was implemented as a set of C++ and Fortran programs (see Data Availability). Manual inspection of results was performed using the FASTA3 program ^53^. The number of mismatches between an insertion sequence (observed or shuffled) and PIS was taken as weight W. The distribution of weights W_shuffled_ was calculated for 1,000 shuffled insertion sequences This distribution was used to calculate probability P(W_observed_ ≥ W_shuffled_). This probability is equal to the number of shuffled insertion sequences with W_shuffled_ equal to or smaller than W_observed_. Small probability values (P(W_observed_ ≥ W_shuffled_) ≤ 0.05) indicate statistical support for the hypothesis that the analyzed insertion sequence results from a duplication or a template switch.

Short inserts (< 9 nucleotides) were marked as a duplication if the insert was identical either to the left or the right adjoining sequence. Additionally, we separated single-nucleotide inserts occurring within homopolymer runs from the rest of the inserts.

### Analysis of transcriptome data and genomic RNA structure

To compare insert locations with RNA secondary structure, we utilized the data from Huston et al., 2021 uploaded to github: https://github.com/pylelab/SARS-CoV-2_SHAPE_MaP_structure. For our analysis we used the data from full-length secondary structure map (.ct file). We considered all paired bases to be in stems, whereas those that are not paired were considered to be located in the loops. Thus, an insert was assigned to the stem if it was flanked on both sides by residues known to be paired.

The data on the SARS-CoV-2 transcriptome was extracted from Ref. 26. Pearson correlation coefficient between insertion locations and template switch hotspots was calculated for bins of size 100 nucleotides with cor.test() function in R version 3.6.3.

To calculate the random distributions for the analyses of distances to the closest junction and appearance of insertions in stems, we performed 1000 permutations, where each time we selected randomly from genome the same number of genome positions as in inserts dataset (347 when we analyzed all inserts, and 45 when we analyzed highly confident inserts). To compare distribution of distances for real data and random data, the Wilcoxon sum rank test was performed. In the case of inserts in stems, the p-value is the portion of cases in our simulation that had the same or smaller number of junctions as the real data.

To analyze whether long insertions coincide with template switch hotspots, we utilized the data on 5’ and 3’ ends of junctions from ^26^. All junction reads within 100 nucleotides from insertion site and insertion source positions were selected. Although the core elements that are involved in template switching are 6-7 nt long, the neighboring regions appear to contribute as well ^23,54^. Furthermore, the peak areas representing the hotspots in the original data are wide, and in the initial publication, 100 nt windows were used for the analysis (Fig. 1e) ^26^. To verify the significance of these findings we performed two simulations. In first scenario the positions of inserts were fixed to the real positions from the data, but the locations of source sequences were randomly sampled 1000 times from the genome, in second scenario both source and insertion site positions were randomly sampled 1000 times. The p-value is the portion of cases in our simulation that have the same or larger number of junctions as the real data.

### Phylogenetic analysis

To find the location of the selected SARS-CoV-2 genomes on the phylogenetic tree we utilized UShER ^55^. The phylogenetic tree of 2,501,152 genomes from GISAID, Genbank, COG-UK and CNCB (2021-07-16) available at UCSC was used as the starting tree, and only leaves representing records from GISAID (1347414 leaves) retained.

To assign PANGO lineages to genomes, we utilized pangolin software v. 3.1.5 ^56^ with default parameters.

An insert was defined as strictly monophyletic if it was observed in at least two genomes, and those genomes formed a stable clade in the phylogenetic tree or were located in the same stem cluster (set of identical sequences in the tree with branches of zero length). Also. we used more relaxed criteria, which implies that inserts in clades that contained less than 1.5% of all genomes in the dataset (2000 leaves) and belonged to the same PANGO lineage were likely monophyletic.

The clades containing the genomes of interest were extracted and vizualized with ETE 3 package for Python ^57^.

### Models of spike protein and visualization

Models were build with SWISS-model ^58^. We used the basic parameters. The models shown on Figure 3 are based on two different initial PDB structures: Cryo-EM structure of PCoV_GX spike glycoprotein (PDB ID: 7cn8), and complex of SARS-CoV-2 spike glycoprotein with 4A8 antibody (PDB ID: 7cl2). The first structure was selected, because it was the best whole length structure with highest aa identity, that cover most of the protein.

The obtained protein models were visualized with Open-Source PyMOL version 2.4.

## Data availability

GISAID data used for this research are subject to GISAID’s Terms and Conditions. SARS-CoV-2 genome sequences and metadata are available for download from GISAID EpiCoV™. The acknowledgements to all Originating and Submitting laboratories are provided in the Supplementary Table 5.

Custom R and Python scripts utilized for data analysis and visualization are available on github: https://github.com/garushyants/covid_insertions_paper

## Acknowledgements

The authors are grateful to Koonin group members for useful discussions. We thank Elena Nabieva for suggestions about variant calling pipelines. This study was supported by the Intramural Research Program of the U.S. National Library of Medicine at the National Institutes of Health.

## Authors contributions

IBR and EVK initiated the study. EVK designed and supervised the project. SKG and IBR collected the data. SKG extracted and verified the inserts, analyzed the data and built protein models. IBR and GSK analyzed the insertion mechanisms and the origins of inserts. GSK and EVK wrote the manuscript that was edited and approved by all authors.

## References

1. Candido, D. S. et al. Evolution and epidemic spread of SARS-CoV-2 in Brazil. Science 369, 1255–1260 (2020).

2. du Plessis, L. et al. Establishment and lineage dynamics of the SARS-CoV-2 epidemic in the UK. Science 371, 708–712 (2021).

3. Munnink, B. B. O. et al. Jumping back and forth: anthropozoonotic and zoonotic transmission of SARS-CoV-2 on mink farms. bioRxiv 2020.09.01.277152 (2020) doi:10.1101/2020.09.01.277152.

4. Komissarov, A. B. et al. Genomic epidemiology of the early stages of the SARS-CoV-2 outbreak in Russia. Nat. Commun. 12, 649 (2021).

5. Martin, D. P. et al. The emergence and ongoing convergent evolution of the N501Y lineages coincides with a major global shift in the SARS-CoV-2 selective landscape. medRxiv 2021.02.23.21252268 (2021) doi:10.1101/2021.02.23.21252268.

6. Davies, N. G. et al. Estimated transmissibility and impact of SARS-CoV-2 lineage B.1.1.7 in England. Science (2021) doi:10.1126/science.abg3055.

7. Tegally, H. et al. Emergence and rapid spread of a new severe acute respiratory syndrome-related coronavirus 2 (SARS-CoV-2) lineage with multiple spike mutations in South Africa. http://medrxiv.org/lookup/doi/10.1101/2020.12.21.20248640 (2020) doi:10.1101/2020.12.21.20248640.

8. Sabino, E. C. et al. Resurgence of COVID-19 in Manaus, Brazil, despite high seroprevalence. The Lancet 397, 452–455 (2021).

9. Planas, D. et al. Reduced sensitivity of SARS-CoV-2 variant Delta to antibody neutralization. Nature 1–7 (2021) doi:10.1038/s41586-021-03777-9.

10. McCarthy, K. R. et al. Recurrent deletions in the SARS-CoV-2 spike glycoprotein drive antibody escape. Science 371, 1139–1142 (2021).

11. Gussow, A. B. et al. Genomic determinants of pathogenicity in SARS-CoV-2 and other human coronaviruses. Proc. Natl. Acad. Sci. 117, 15193–15199 (2020).

12. Andersen, K. G., Rambaut, A., Lipkin, W. I., Holmes, E. C. & Garry, R. F. The proximal origin of SARS-CoV-2. Nat. Med. 26, 450–452 (2020).

13. Walls, A. C. et al. Structure, Function, and Antigenicity of the SARS-CoV-2 Spike Glycoprotein. Cell 181, 281-292.e6 (2020).

14. Peacock, T. P. et al. The furin cleavage site in the SARS-CoV-2 spike protein is required for transmission in ferrets. Nat. Microbiol. 6, 899–909 (2021).

15. Singer, J., Gifford, R., Cotten, M. & Robertson, D. CoV-GLUE: A Web Application for Tracking SARS-CoV-2 Genomic Variation. (2020) doi:10.20944/preprints202006.0225.v1.

16. Chrisman, B. S. et al. Indels in SARS-CoV-2 occur at template-switching hotspots. BioData Min. 14, 20 (2021).

17. Li, X. et al. Emergence of SARS-CoV-2 through recombination and strong purifying selection. Sci. Adv. 6, (2020).

18. Boni, M. F. et al. Evolutionary origins of the SARS-CoV-2 sarbecovirus lineage responsible for the COVID-19 pandemic. Nat. Microbiol. 5, 1408–1417 (2020).

19. Graham, R. L. & Baric, R. S. Recombination, Reservoirs, and the Modular Spike: Mechanisms of Coronavirus Cross-Species Transmission. J. Virol. 84, 3134–3146 (2010).

20. Xiao, K. et al. Isolation of SARS-CoV-2-related coronavirus from Malayan pangolins. Nature 583, 286–289 (2020).

21. Simon-Loriere, E. & Holmes, E. C. Why do RNA viruses recombine? Nat. Rev. Microbiol. 9, 617–626 (2011).

22. Sethna, P. B., Hung, S. L. & Brian, D. A. Coronavirus subgenomic minus-strand RNAs and the potential for mRNA replicons. Proc. Natl. Acad. Sci. 86, 5626–5630 (1989).

23. Sawicki, S. G., Sawicki, D. L. & Siddell, S. G. A Contemporary View of Coronavirus Transcription. J. Virol. 81, 20–29 (2007).

24. V’kovski, P., Kratzel, A., Steiner, S., Stalder, H. & Thiel, V. Coronavirus biology and replication: implications for SARS-CoV-2. Nat. Rev. Microbiol. 19, 155–170 (2021).

25. Nomburg, J., Meyerson, M. & DeCaprio, J. A. Pervasive generation of non-canonical subgenomic RNAs by SARS-CoV-2. Genome Med. 12, 108 (2020).

26. Kim, D. et al. The Architecture of SARS-CoV-2 Transcriptome. Cell 181, 914-921.e10 (2020).

27. Quéromès, G. et al. Characterization of SARS-CoV-2 ORF6 deletion variants detected in a nosocomial cluster during routine genomic surveillance, Lyon, France. Emerg. Microbes Infect. 10, 167–177 (2021).

28. Huston, N. C. et al. Comprehensive in vivo secondary structure of the SARS-CoV-2 genome reveals novel regulatory motifs and mechanisms. Mol. Cell 81, 584-598.e5 (2021).

29. Kondrashov, A. S. & Rogozin, I. B. Context of deletions and insertions in human coding sequences. Hum. Mutat. 23, 177–185 (2004).

30. Hausmann, S., Garcin, D., Delenda, C. & Kolakofsky, D. The versatility of paramyxovirus RNA polymerase stuttering. J. Virol. 73, 5568–5576 (1999).

31. Zheng, H., Lee, H. A., Palese, P. & García-Sastre, A. Influenza A Virus RNA Polymerase Has the Ability To Stutter at the Polyadenylation Site of a Viral RNA Template during RNA Replication. J. Virol. 73, 5240–5243 (1999).

32. Pfeiffer, F. et al. Systematic evaluation of error rates and causes in short samples in next-generation sequencing. Sci. Rep. 8, 10950 (2018).

33. Rang, F. J., Kloosterman, W. P. & de Ridder, J. From squiggle to basepair: computational approaches for improving nanopore sequencing read accuracy. Genome Biol. 19, 90 (2018).

34. Ma, X. et al. Analysis of error profiles in deep next-generation sequencing data. Genome Biol. 20, 50 (2019).

35. Dohm, J. C., Peters, P., Stralis-Pavese, N. & Himmelbauer, H. Benchmarking of long-read correction methods. NAR Genomics Bioinforma. 2, (2020).

36. te Velthuis, A. J. W., Arnold, J. J., Cameron, C. E., van den Worm, S. H. E. & Snijder, E. J. The RNA polymerase activity of SARS-coronavirus nsp12 is primer dependent. Nucleic Acids Res. 38, 203–214 (2010).

37. Ferron, F. et al. Structural and molecular basis of mismatch correction and ribavirin excision from coronavirus RNA. Proc. Natl. Acad. Sci. 115, E162–E171 (2018).

38. Rochman, N. D. et al. Ongoing global and regional adaptive evolution of SARS-CoV-2. Proc. Natl. Acad. Sci. U. S. A. 118, e2104241118 (2021).

39. Davidson, A. D. et al. Characterisation of the transcriptome and proteome of SARS-CoV-2 reveals a cell passage induced in-frame deletion of the furin-like cleavage site from the spike glycoprotein. Genome Med. 12, 68 (2020).

40. Kemp, S. A. et al. SARS-CoV-2 evolution during treatment of chronic infection. Nature 1–10 (2021) doi:10.1038/s41586-021-03291-y.

41. Sepulcri, C. et al. The longest persistence of viable SARS-CoV-2 with recurrence of viremia and relapsing symptomatic COVID-19 in an immunocompromised patient – a case study. medRxiv 2021.01.23.21249554 (2021) doi:10.1101/2021.01.23.21249554.

42. Cerutti, G. et al. Potent SARS-CoV-2 neutralizing antibodies directed against spike N-terminal domain target a single supersite. Cell Host Microbe 29, 819-833.e7 (2021).

43. Tarke, A. et al. Comprehensive analysis of T cell immunodominance and immunoprevalence of SARS-CoV-2 epitopes in COVID-19 cases. bioRxiv 2020.12.08.416750 (2020) doi:10.1101/2020.12.08.416750.

44. Rosa, A. et al. SARS-CoV-2 can recruit a haem metabolite to evade antibody immunity. Sci. Adv. eabg7607 (2021) doi:10.1126/sciadv.abg7607.

45. Johnson, B. A. et al. Loss of furin cleavage site attenuates SARS-CoV-2 pathogenesis. Nature 1–7 (2021) doi:10.1038/s41586-021-03237-4.

46. Papa, G. et al. Furin cleavage of SARS-CoV-2 Spike promotes but is not essential for infection and cell-cell fusion. PLOS Pathog. 17, e1009246 (2021).

47. Kim, M.-J. & Kao, C. Factors regulating template switch in vitro by viral RNA-dependent RNA polymerases: Implications for RNA–RNA recombination. Proc. Natl. Acad. Sci. 98, 4972–4977 (2001).

48. Simon-Loriere, E. & Holmes, E. C. Why do RNA viruses recombine? Nat. Rev. Microbiol. 9, 617–626 (2011).

49. Yang, Y., Yan, W., Hall, A. B. & Jiang, X. Characterizing Transcriptional Regulatory Sequences in Coronaviruses and Their Role in Recombination. Mol. Biol. Evol. 38, 1241–1248 (2021).

50. MacLean, O. A. et al. Natural selection in the evolution of SARS-CoV-2 in bats created a generalist virus and highly capable human pathogen. PLoS Biol. 19, e3001115 (2021).

51. Langmead, B. & Salzberg, S. L. Fast gapped-read alignment with Bowtie 2. Nat. Methods 9, 357–359 (2012).

52. Wilm, A. et al. LoFreq: a sequence-quality aware, ultra-sensitive variant caller for uncovering cell-population heterogeneity from high-throughput sequencing datasets. Nucleic Acids Res. 40, 11189–11201 (2012).

53. Pearson, W. R. Finding Protein and Nucleotide Similarities with FASTA. Curr. Protoc. Bioinforma. 53, 3.9.1-3.925 (2016).

54. Sola, I., Almazán, F., Zúñiga, S. & Enjuanes, L. Continuous and Discontinuous RNA Synthesis in Coronaviruses. Annu. Rev. Virol. 2, 265–288 (2015).

55. Turakhia, Y. et al. Ultrafast Sample placement on Existing tRees (UShER) enables real-time phylogenetics for the SARS-CoV-2 pandemic. Nat. Genet. 53, 809–816 (2021).

56. Rambaut, A. et al. A dynamic nomenclature proposal for SARS-CoV-2 lineages to assist genomic epidemiology. Nat. Microbiol. 5, 1403–1407 (2020).

57. Huerta-Cepas, J., Serra, F. & Bork, P. ETE 3: Reconstruction, Analysis, and Visualization of Phylogenomic Data. Mol. Biol. Evol. 33, 1635–1638 (2016).

58. Waterhouse, A. et al. SWISS-MODEL: homology modelling of protein structures and complexes. Nucleic Acids Res. 46, W296–W303 (2018).

