## Supplemental information for "Insertions in SARS-CoV-2 genome caused by template switch and duplications give rise to new variants that merit monitoring"

### Supplementary Information

#### Supplementary Tables

**Supplementary Table 1. The list of inserts in SARS-CoV-2 genomes and genomes where they were found.** Abbreviations in LoFreq Info column: DP – Raw Depth, AF – Allele Frequency, SB – Phred-scaled strand bias at this position, DP4 – Counts for ref-forward bases, ref-reverse, alt-forward and alt-reverse bases, HRUN – Homopolymer length to the right of report indel position.

**Supplementary Table 2.** The summary of supporting data for 354 inserts observed in this study. The details on the monophyly test are provided in Materials and Methods.

**Supplementary Table 3.** SRA records information for genomes found on the same clade as sequences containing ins22204:ATAGATCGA and ins22205:CGGCAGGCT.

**Supplementary Table 4.** The list of high confidence inserts with the potential mechanisms of their appearance. For template switch candidate the location of origin, sequence in origin and p-value are specified.

**Supplementary Table 5.** The acknowledgments to all Originating and Submitting laboratories that provided the data to GISAID utilized in this paper.

#### Supplementary Notes

##### Supplementary Note 1

##### SARS-CoV-2 phylogeny and origin of inserts

Insertion, especially long ones, are assumed to be rare events, and therefore, all inserts in the same genomic position are generally expected to be monophyletic. To test this prediction, we mapped genomes containing inserts on the SARS-CoV-2 tree constructed with UShER (see Materials and Methods). Some inserts indeed were found to be monophyletic, but in other cases, genomes with inserts appear close on the tree but not strictly monophyletic, being interspersed with genome lacking the respective insertion (Supplementary Figure 2,3).

In principle, there could be three main causes behind this unexpected pattern: 1) inserts are indeed not monophyletic, that is, insertion occurred in the same position on several independent occasions, 2) the genomes were incorrectly placed in the tree, 3) inserts are not always called, such that some genomes appear to be lacking an insert that, in actuality, they contain. To discriminate between these possibilities, we investigated the non-singleton long inserts in the S gene in more detail.

To rule out the erroneous phylogeny hypothesis, we rebuilt the phylogenetic tree with IQtree2 for the respective individual clades. In all cases, the topology was similar to the one produced by UShER, which suggests that this topology is not an error of the phylogeny reconstruction.

We then searched for available raw sequencing data for the genomes that lacked insertion in a given position, while belonging to the same clade with genomes containing the insert. For two cases (ins22204:ATAGATCGA and ins22205:CGGCAGGCT), we detected 4 and 38 records in SRA, respectively (Supplementary table 3).

In all four cases, the raw read data supported ins22204. Ins22205 insertion was supported in 28 cases, in 6 others, the region containing the insertion was either not covered of covered by less than 10 reads, which included the insert, in 2 more cases, the insert was present, but in < 50% of the corresponding reads, and in only 2 cases, the insertion was not supported by the raw read data. The genomes with supported inserts were scattered across the tree (Supplementary Figure 2,3), suggesting that inserts were not called in a large fraction of the genomes in the respective clades.

##### Supplementary note 2

Among eight high confidence inserts observed in our dataset, only one, ins246DSWG, was observed in a VOC (B.1.1.7 or Alpha). This insert is located within the epitope interacting with antibodies in NTD and thus merits close monitoring.

Other inserts occurred in less prominent lineages, such as A.2.5, B.1.1, B.1.177, B.1.214.2, B.1.1.372, B.1.596, and B.1.1.70 (Supplementary table 4). While lineages B.1.177, B.1.1.372, B.1.596 and B.1.1.70 potentially included both genomes with and without the inserts and were almost completely displaced, presumably by the Delta variant by July 2021 (<https://outbreak.info/>), lineages A.2.5 (ins214AAG), B.1.214.2 (ins214TDR) started to grow in frequency in the beginning of 2021 and are still observed in population. Thus, these might be antibody escape variants that are being selected in response to vaccination.

The entire B.1.214.2 and A.2.5 PANGO lineages are the same clades that contain inserts ins214TDR (ins22204) and ins214AAG (ins22205), respectively (Supplementary note 1). Thus, the inserts are previously undetected signature features of these PANGO lineages. We further investigated the temporal dynamics of these two inserts by plotting the number of genomes in the respective clades (Supplementary note 1) collected on a particular date. We confirmed the presence of these inserts in our data in early January 2021, and show that they are present at least till the end of the observation period, e.g. the beginning of July 2021 (Supplementary Figure 2b,3b). The appearance of these lineages contemporaneously with the start of mass vaccination together with their location on the protein surface, suggests that these two inserts might lead to partial antibody escape. Furthermore, lineage A.2.5 mostly includes samples from Latin America ^1^, whereas B.1.214.2 is predominant in Kongo ^2^. Both these regions are substantially under-sampled compared to North America and Europe, so that the frequencies of these variants are likely to be underestimated, that is why we propose that ins214TDR and ins214AAG require further monitoring.

**References**

1. A.2.5 Lineage Report. *outbreak.info* https://outbreak.info/situation-reports?pango=A.2.5&loc=USA&loc=USA_US-CA&selected=Worldwide.

2. B.1.214.2 Lineage Report. *outbreak.info* https://outbreak.info/situation-reports?pango=B.1.214.2&loc=USA&loc=USA_US-CA&selected=Worldwide.

#### Supplementary Figures

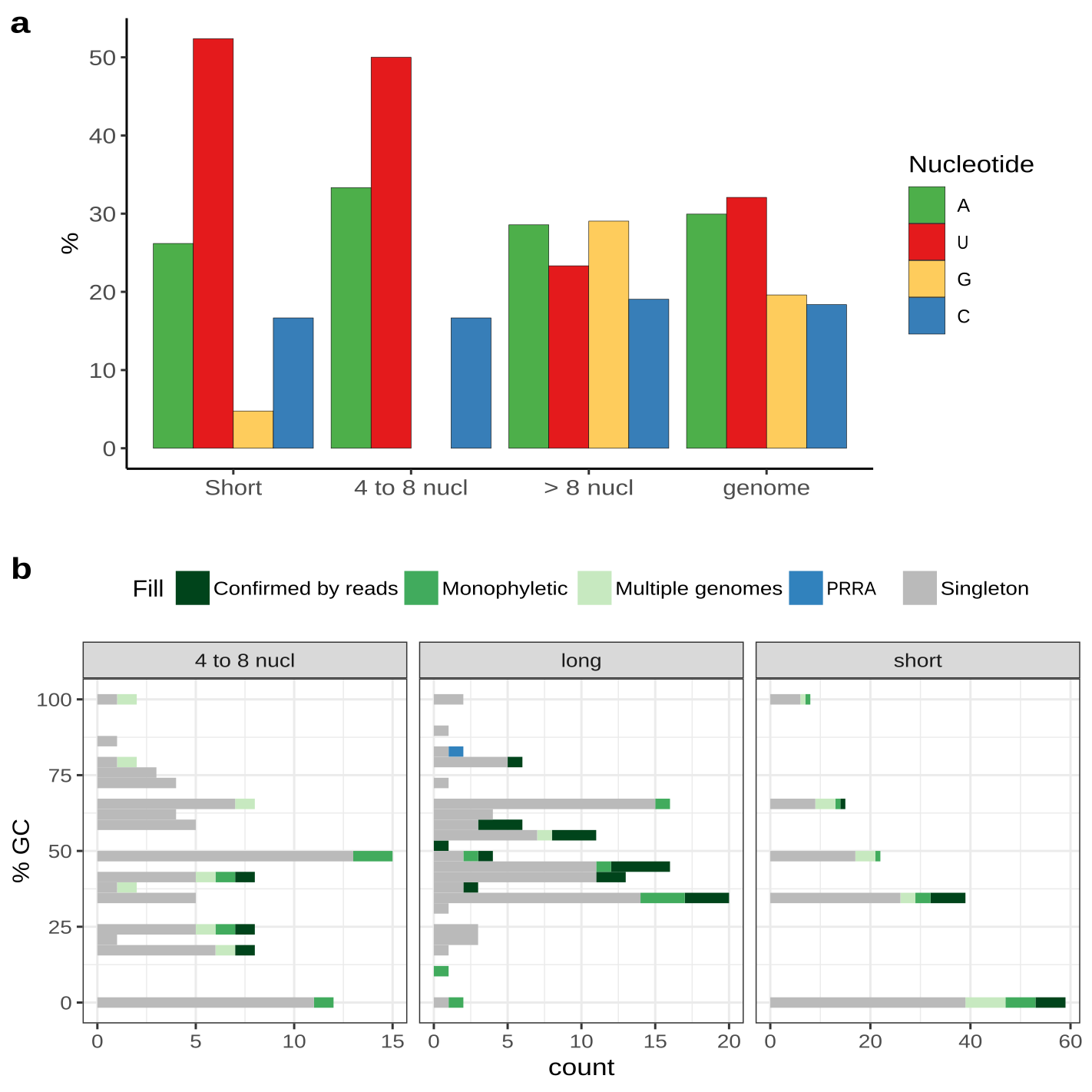
Supplementary Figure 1. **Nucleotide composition of the insert**s.

(a) Nucleotide composition of inserts is shown as in Figure 1b, but only for inserts that were validated by sequencing data analysis. (b) GC-content of in inserts of different length. Shown in blue is the PRRA insert that introduces a polybasic furin cleavage site into the S protein, and is one of the characteristic features of SARS-CoV-2.

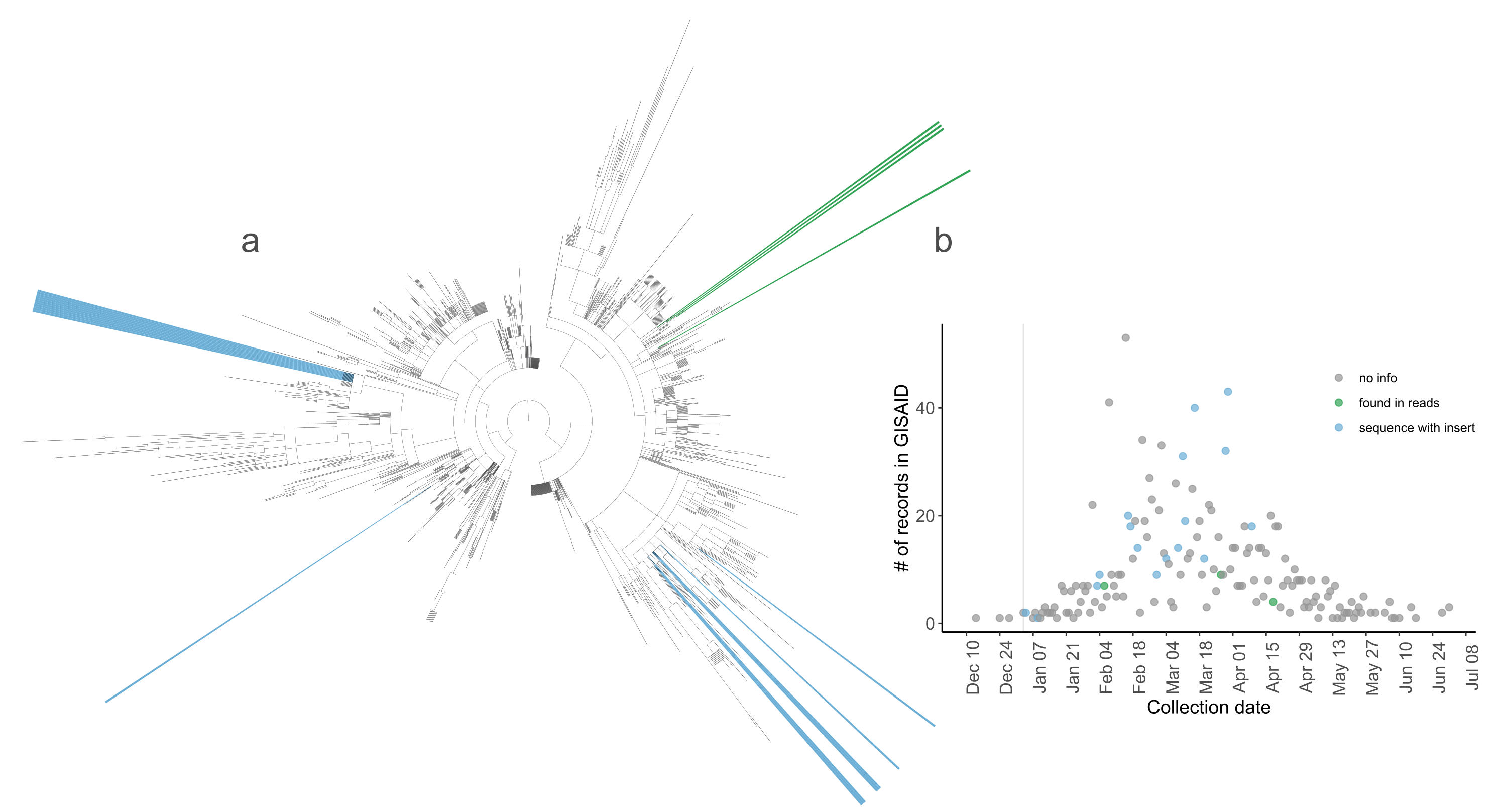

Supplementary Figure 2. **Evolutionary history of ins22204**.

(a) Phylogenetic tree for the clade containing ins22204:ATAGATCGA. (b) Number of samples from the clade collected at particular date. Grey vertical line shows the earliest date when we were able to confirm the insert.

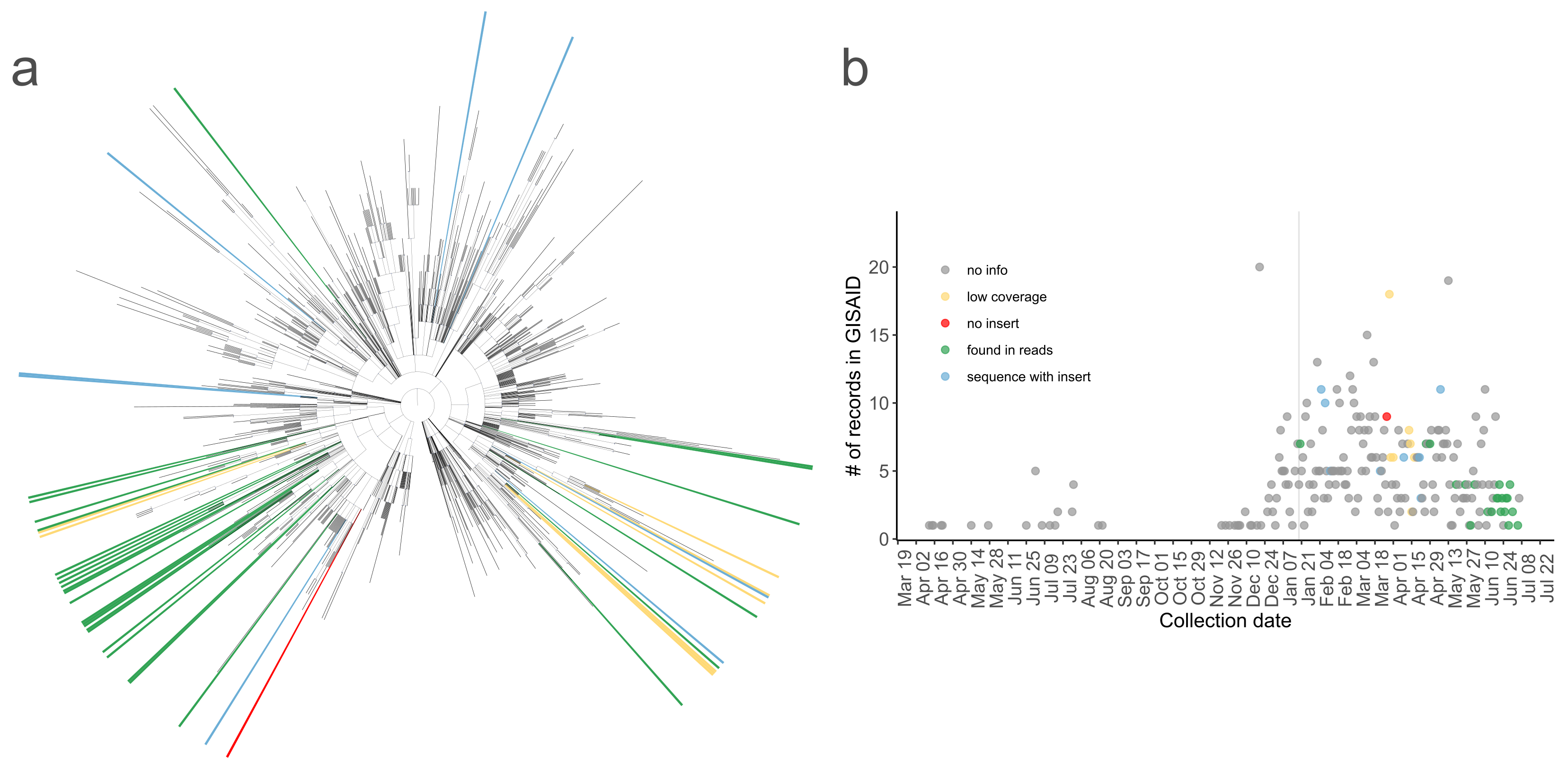

Supplementary Figure 3. **Evolutionary history of ins22205.**

(a) Phylogenetic tree for the clade containing ins22205:CGGCAGGCT. (b) Number of samples from the clade collected at particular date. Grey vertical line shows the earliest date when we were able to confirm the insert.

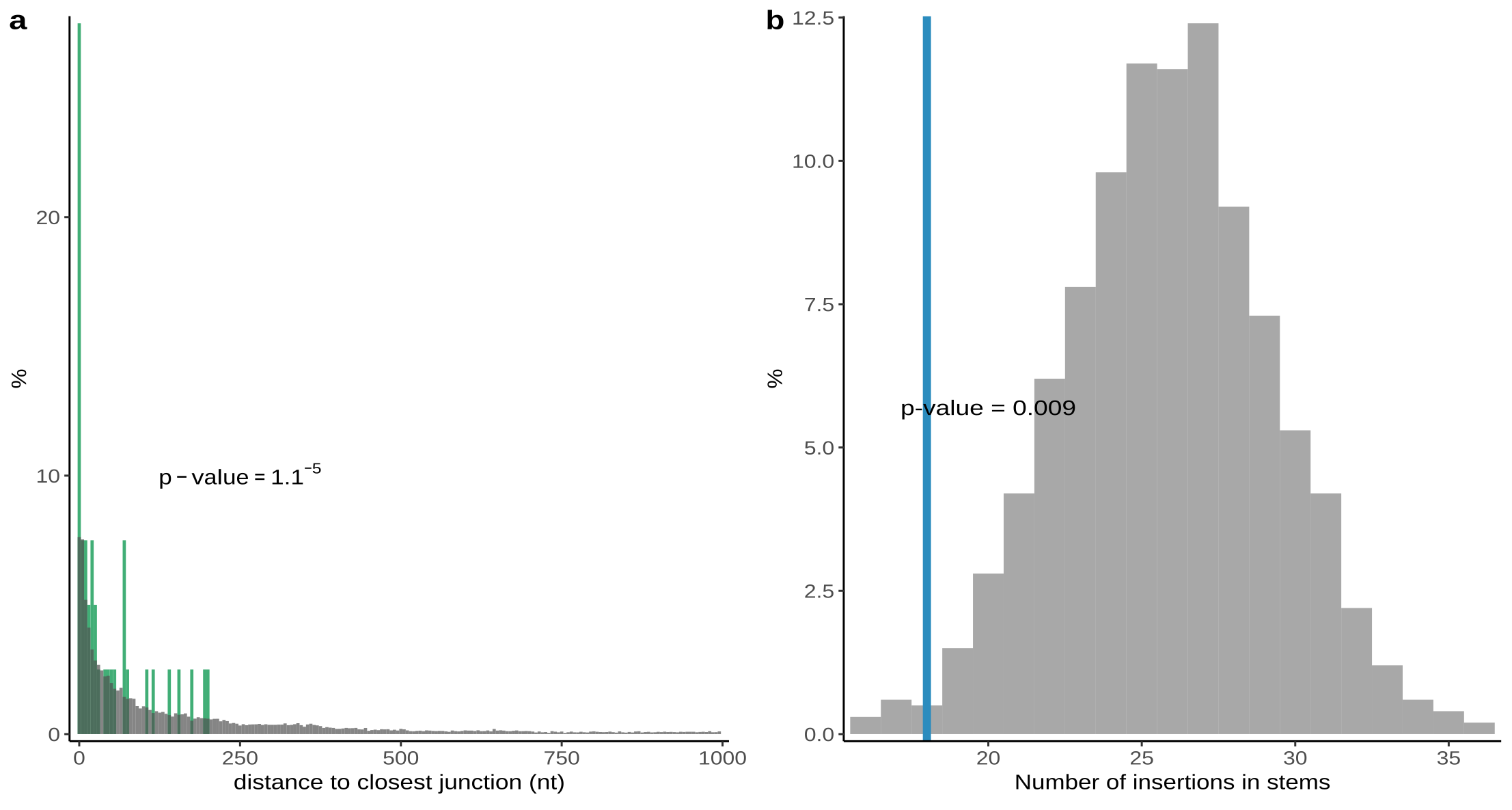

Supplementary Figure 4. **Patterns of insert distribution across SARS-CoV-2 genomes.**

(a) Distance from highly confident inserts to closest template switch hotspot site (green) in comparison with random expectation (grey).

(b) The number of highly confident inserts that occur in structured regions of SARS-CoV-2 genomic RNA (blue) in comparison with random expectation (grey). The data on SARS-CoV-2 structure is obtained from Huston et al., 2021. The inserts identified with high confidence include 17 short insertions confirmed by raw read analysis, and 28 long inserts that were either confirmed by raw read analysis or found in multiple genomes and shown to be monophyletic.

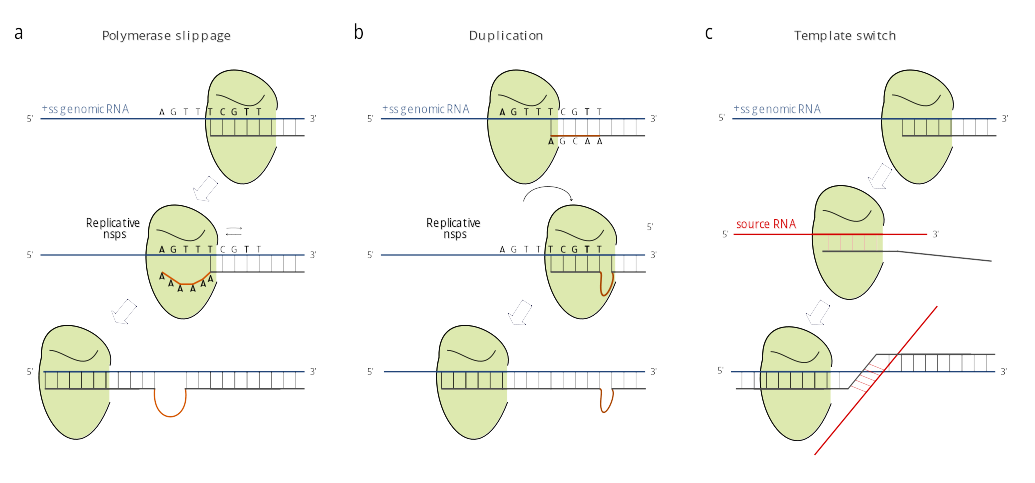

Supplementary Figure 5. **Possible mechanisms of insertions**

(a) polymerase slippage

(b) local duplication

(c) template switch.
